# Desire and Craving ratings vary significantly for healthy alcohol consumers: Differences in semantic interpretation?

**DOI:** 10.1101/2021.10.25.465749

**Authors:** H Peterson, LJ Veach, SL Simpson, J Fanning, PJ Laurienti, L Gauvin

## Abstract

Craving is a central concept in alcohol, and other substance, research. Beginning in 1955 the World Health Organization outlined a working definition of the term to be used in research and clinical settings. However, the semantic interpretation of “craving” as a concept is not widely agreed upon. Since the publication of this first craving definition, a handful of studies have been conducted to investigate differences in operational definitions of “craving”, and have demonstrated a lack of agreement between studies and across research subjects. With this background as evidence, our research group investigated, when left to their own semantic understanding of the terms, if regular alcohol consumers would rate craving for alcohol and desire for alcohol in similar ways using related descriptors. Thirty-nine healthy, non-binging regular alcohol consumers were studied across periods of their typical alcohol consumption and imposed alcohol abstinence, collecting ratings of desire and craving for alcohol approximately every two hours across the two experimental periods, and during neutral and alcohol related imagery viewing. Among these non-binging regular drinkers, ratings of desire and craving for alcohol are consistently different while drinking according to a person’s typical routine or abstaining, throughout the day, and when viewing alcohol cue imagery.

## INTRODUCTION

Millions of people worldwide consume alcohol: upwards of 50% of the US population reports alcohol consumption within the last month (SAMHSA, 2015, Collaborators, 2018). Along with these high rates of consumption, National Institutes of Health (NIH) studies have begun to demonstrate increased prevalence of riskier use that does not meet alcohol use disorder (AUD) diagnostic criteria (e.g. had times when they drank more or longer than intended, wanted to cut down or stop drinking, found that drinking interfered with taking care of home or family, or continued drinking even though it is causing trouble in personal relationships), in addition to mild, moderate or severe AUD (White et al., 2011, Grant et al., 2015, NIAAA, 2015, NIH and DHHS, 2015, 2017, Higson et al., 2017, Blow et al., 2020, Riggs et al., 2020). Today, a wide body of research has documented the important role craving plays in alcohol consumption: craving is a driving factor of alcohol consumption (Rankin et al., 1979, Ludwig, 1986, Kozlowski et al., 1989, de Wit, 2000, May et al., 2014, Mayhugh et al., 2018a, Mayhugh et al., 2018c), one diagnostic feature of AUD (APA, 2013), a symptom of withdrawal (Sullivan et al., 1989b), and a key predictor of relapse in patients with AUD (Breese et al., 2006, Gordon et al., 2006, Moos and Moos, 2006, Sinha, 2007, Miller and Gold, 2011, Sinha, 2011, Sinha, 2012, Seo et al., 2013, Volkow and Baler, 2013, Tuithof et al., 2014, Muller and Meyerhoff, 2020, Weiss et al., 2020).

As such, “craving” is a central concept of alcohol research. In 1955, the World Health Organization convened to define craving as a central part of the alcoholic experience (W.H.O., 1955). At the time, vernacular use of the term typically conveyed “urgent and overpowering desire”, an “irresistible impulse”, or “a strong desire or intense longing” (Fowler and Fowler, 1964, Kozlowski and Wilkinson, 1987). Since this preliminary interest in the study of alcohol craving, operational definitions have grown to vary across studies (W.H.O., 1955, Kozlowski et al., 1989, Breese et al., 2006, Gordon et al., 2006, Miller and Gold, 2011, Tuithof et al., 2014). As early as 1979, researchers began to investigate the potential differences between (1) a “craving” (a yearning for the *effects* of alcohol) versus (2) simply a desire to consume alcohol, and the physiological differences between these states (Rankin et al., 1979). Self-reported differences were found between ratings of “desire for a drink” and “difficulty in resisting alcohol” in persons with AUD (Rankin et al., 1979). This lack of agreement is not confined to alcohol cravings: in a study of cigarette smokers undergoing alcohol or drug treatment at the Clinical Institute of the Addiction Research Foundation, the most common response to “which best describes what *you* meant by a *craving* for a drug” was “a *strong* urge or desire to take a drug” (p. 443) (Kozlowski et al., 1989). Nevertheless, greater than 35% of the sample said “craving” was in fact “*any* urge or desire”, and 15% said neither of these definitions fit their understanding of the term (p. 443) (Kozlowski et al., 1989). Two thirds of a population of individuals with an AUD gave the same rating for a “desire” and a “craving” for an alcoholic drink, while a quarter rated “craving” lower than “desire”, and a small portion of the sample even rated “craving” higher than “desire” for a drink (Kozlowski et al., 1989). Even these early studies demonstrate a lack of agreement between both research studies and research subjects as to the interpretation or understanding of “craving”.

When considering the clinical significance of “craving”, definitions range from “a subjective experience of wanting to use a drug” (p.24) (Tiffany and Wray, 2012) to “an intense, conscious desire, usually to consume a specific drug or food” (p. 101) (Tapper, 2018) or “a powerful subjective experience that motivates [drug seeking]” (p. 1061) (Caselli and Spada, 2011). Many questionnaires that assess craving and desire are available, including the Craving Experience Questionnaire (May et al., 2014), the Desires for Alcohol Questionnaire (Pasche et al., 2013) and Desire Thinking Questionnaires (Kavanagh et al., 2005, Kavanagh et al., 2009, Caselli and Spada, 2011). Across these three surveys alone, questions include the terms “want”, “need”, “urge”, “think”, and “intrusive feelings” (May et al., 2014). Are these terms synonymous with “craving” for respondents, or does “craving” encapsulate these multiple psychological experiences? The Desires for Alcohol Questionnaire is comprised of questions focused entirely on “drinking” (e.g. “drinking right now would make me feel less tense”), with only one of fourteen items probing “desire to drink” (e.g. “my desire to drink right now seems overwhelming”) (Pasche et al., 2013), demonstrating the wide and conflicting, at times, range of operational definitions of “craving” even among this small number of quantitative data collection tools. Exploring differences, if any, between craving and desire may also serve to provide more clinical tools as drinkers examine their patterns of alcohol use.

With this background as evidence, our research group investigated if regular alcohol consumers would rate craving for alcohol and desire for alcohol in similar ways using related descriptors. As we unravel the important neurobiological differences between drinking and abstinence experiences (Mayhugh et al., 2018a, Mayhugh et al., 2018c, Peterson et al., Under Review-c), we hope to accurately capture the unique constructs of craving and desire as risk factors for developing AUD, as indicators of relapse risk, and as motivators for the drinker seeking clinical assessments. Does probing alcohol craving fully capture a consumer’s drive to drink? Or does craving apply as a descriptor to multiple physiological and psychological states? These questions apply to a wide range of research areas, such as alcohol use with obsessive compulsive disorder (OCD), food, and nicotine (Castellani and Rugle, 1995, Kranzler et al., 1999, Anton, 2000, May et al., 2004, Field et al., 2008, Moreno et al., 2009, Caselli and Spada, 2011, Lievaart et al., 2015), thus demonstrating the breadth of the need to more fully understand how research participants describe, define, and experience “craving”, if different from desire.

Specifically, potential differences between ratings of alcohol “craving” and “desire” are the primary focus in this examination. Healthy, non-binging daily alcohol consumers were studied, as they make up a large proportion of the current drinking population (NIAAA, 2011, NSDUH, 2019). Additionally, previous studies in this population from our laboratory (unpublished) anecdotally showed that these regular consumers (Mayhugh et al., 2018a, Mayhugh et al., 2018c, Peterson et al., Under Review-c) viewed “craving” as too clinical a term to describe their “desire” to have a drink. As evidence of this report, these regular drinkers reported very low levels of craving, even following stress during a period of abstinence (Mayhugh et al., 2018c). Therefore, this follow-up study set out to examine potential differences in self-reported ratings for “desire” and “craving” for alcohol in regular drinkers. Of note, this study attempted to examine non-AUD alcohol consumers, however participants in this study were not assessed by a clinician or licensed addiction specialist. Instead, the study team was careful to exclude individuals with a known AUD diagnosis and potential participants who indicated experiencing potential AUD symptoms as assessed by DSM-V questions. In this study, craving and desire ratings were probed throughout the day and following alcohol cue exposure, both during typical alcohol consumption and imposed abstinence, in an effort to widely capture potential differences in ratings.

## METHODS AND MATERIALS

### Study Overview

The data presented in the following analyses come from a multi-visit study examining neurobiological variables in healthy, moderate to heavy alcohol consumers (Figure 1). The study protocol included an initial screening visit to determine participant eligibility and collect informed consent followed by a behavioral baseline visit. Participants were assigned to complete 3-day experimental periods of their typical alcohol consumption (typical drinking) and of complete imposed abstinence (abstinence) with the order randomized across participants. During both experimental periods, participants completed surveys on study-issued mobile devices approximately every two hours, following an Ecological Momentary Assessment (EMA) protocol (Stone and Shiffman, 1994, Moskowitz and Young, 2006, Shiffman et al., 2008). Following each experimental period (on the fourth day), participants underwent magnetic resonance imaging (MRI) scanning, during which functional data were collected while participants viewed neutral and alcohol-related images. The current manuscript focuses on craving and desire ratings collected in the EMA surveys as well as ratings collected following the functional MRI scans.

**Figure 1.**
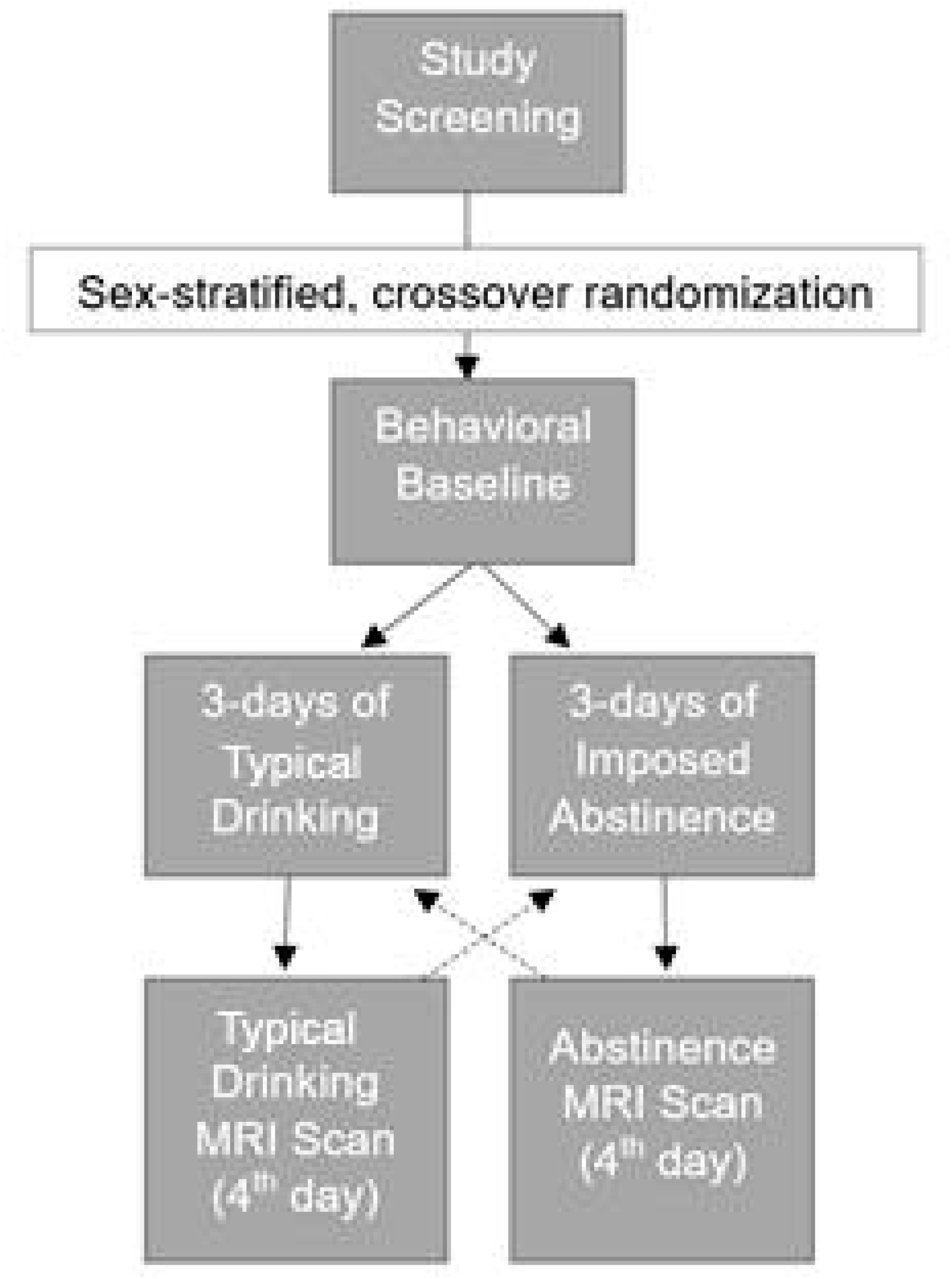
Full study protocol; participants completed four in-person study visits, with three consecutive days of their typical drinking routine or alcohol abstinence prior to each MRI visit. Order of the experimental trials was randomized in a sex-stratified crossover design.

### Participants

Participants (n = 39) were recruited from the Winston-Salem, NC community (22 females). Inclusion criteria included: age between 24-60 years, right-handed, self-reported consumption of alcohol on at least 50% of days (over the previous 90 days), and average drinking pattern of at least 7 drinks/week for females and 14 drinks/week for males (to ensure consumption at or above NIAAA low risk levels (NIAAA)) for at least three years. Exclusion criteria included: current or past AUD diagnosis or self-reported problems with drugs or alcohol, binge drinking episodes (as defined by the NIAAA, at least 4 drinks for females and 5 drinks for males consumed in 2 hours) more than once monthly on average, regular morning drinking or “eye openers” (alcohol consumed before noon), current treatment for any serious illnesses, scores greater than 20 on the Center for Epidemiological Studies Depression Scale (CES-D) (Radloff, 1977), diagnosis of any neurological diseases or psychiatric conditions (diagnosed depression was allowed if stably treated for at least 2 months), consumption of more than 500 mg of caffeine daily, smoking more than 1.5 packs/day or equivalent nicotine use, or a positive saliva drug test (monitoring methamphetamine, cocaine, marijuana, amphetamine, opiates, and benzodiazepines). Because previous research has associated body mass index (BMI) with blood-alcohol concentration (Wang et al., 1992), BMI was restricted to at least 18.5 kg/m^2^ and no more than 35 kg/m^2^. The Clinical Institute Withdrawal Assessment of Alcohol, revised (CIWA-Ar) was used to screen all participants for withdrawal symptoms after the experimental period of abstinence (Sullivan et al., 1989a); any participant exhibiting scores above 7 would be excluded on the basis of potential alcohol dependence (although a positive result did not occur). All participants received a total of $350 for complete study participation. After removal of missing data points, full datasets (EMA and cue-viewing ratings) from 32 participants (35 for only EMA responses) are presented here.

### Procedure

Study procedures were scheduled to avoid any major life stressors or atypical routines. All participants initially underwent a telephone screening to determine study eligibility. The study protocol then consisted of four in-person visits: a screening visit, a second behavioral visit, and two magnetic resonance imaging (MRI) visits in a sex-stratified randomized order. During the screening and baseline visits, all participants gave written informed consent approved by the Wake Forest Health Sciences IRB and completed a battery of self-report questionnaires, including the Timeline Followback (TLFB) (Sobell and Sobell, 2000, Vakili et al., 2008), and scales of stress (Cohen et al., 1983), anxiety (Julian, 2011), mindfulness (Reise et al., 2013), trait craving (Statham et al., 2011), and self-control over their drive to drink (Lowe et al., 2009).

Following the first two study visits, prior to each MRI scan, participants completed surveys on mobile devices, using an EMA protocol for 3-days of their typical drinking and 3-days of abstinence. EMA data were collected throughout both experimental trials (Wray et al., 2014), and involved responding to brief, six question surveys upon waking, approximately every 2-hours as prompted via text message, and before going to bed. Each survey probed stress, craving, desire, arousal, affect, and mindfulness. Responses to questions relevant to craving and desire are presented here. These questions probed “How much do you crave a drink right now?” and “How much do you desire a drink right now?” respectively. Responses were provided on a 0 to 1000 numeric rating scale. Analyses of responses to stress and craving in a similar sample have been published in prior manuscripts (Mayhugh et al., 2018a, Mayhugh et al., 2018c), and additional manuscripts are published examining responses to stress, affect, and arousal (Peterson et al., Under Review-a, Peterson et al., Under Review-b).

Following each of the 3-day experimental periods, participants underwent MRI scanning while viewing neutrally rated images from the International Affective Picture Set (IAPS) (Lang et al., 2008a) followed by alcohol-related images from the Geneva Appetitive Alcohol Pictures (GAAP) (Billieux et al., 2011b). Each scan ran for a duration of 7 minutes, during which participants viewed IAPS and GAAP images for 10 seconds each, resulting in viewing of approximately 42 images from each set. Images from the IAPS were considered “neutral” if their average validation ratings ranged from 4.5-5.5 on valence and arousal scales (overall range 1-9) (Lang et al., 2008b), including images depicting, for example, a rolling pin (image 7000), a collection of buttons (image 7001), a lamp (image 7175), and a box of tissues (image 7950). Selected GAAP images were also rated, on average, between 4.5-5.5 on validation scales of valence and arousal (Billieux et al., 2011a), and included images depicting, for example, a corkscrew (image 16), a glass of red wine (image 26), a six-pack of beer bottles (image 46), and a collection of liquor bottles on a bar (image 15).

Following cue viewing, participants were presented a visual analogue scale (VAS) ranging from “not at all” to “extremely” with corresponding questions probing their current craving and desire. Technicians moved a computer cursor across the VAS, and participants verbally chose where to place their selection to avoid unnecessary movement in the scanner. Responses were collected on a 0 – 1000 scale (no numbers shown to participants) to make the selection as smooth (rather than stepped) as possible. All MRI scans were conducted at a large southeastern U.S. academic medical center, scheduled during the time participants would typically be drinking (typically between 5-8 PM). The scanning protocol was the same for both the typical drinking and abstained MRI visits. Brain imaging results will be presented in manuscripts dedicated to the neurobiological hypotheses of the study.

### EMA Statistical Modeling

EMA results discussed in this analysis focused on desire and craving ratings collected during the typical drinking period and during the abstinence period. Results were computed using a 2-level nested hierarchical model on 1,598 EMA responses from 35 participants. All EMA responses were concatenated and tallied to record number of surveys completed. Responses were categorized as typical drinking or abstained responses, and responses from the typical drinking period were stratified as pre-drinking or post-drinking using dummy variables. Time of day was recorded in 24-hour notation, and across participants, time was centered at 15:00 (3 PM). Responses recorded after midnight belonging to the prior day were reassigned times greater than 24:00 to accurately reflect the time of day at which they occurred (i.e. 25:00 for 1 AM).

When creating the statistical model, the centered time (15:00) was squared to operationalize quadratic trends to account for potential nonlinear variations in ratings across the day (Mayhugh et al., 2018a, Mayhugh et al., 2018b). Multivariate modeling was used to determine the influence of within and between-subject variations (Raudenbush, 2001). A null model was applied in order to estimate the intraclass correlation (ICC) and to decompose the variance between and within participants. The variables operationalizing the linear and quadratic trends of time of day were entered into the statistical model to explore within-day variations in ratings, and a variable contrasting drinking states was entered in the model to examine the effect of drinking state (Mayhugh et al., 2018a, Mayhugh et al., 2018b). All EMA analyses were conducted using HLM 8 software (Raudenbush et al., 2019).

### In-Scanner Response Analysis

Craving and desire ratings collected in the MRI scanner following the neutral imagery and alcohol imagery tasks were compared across tasks and across experimental conditions. The distribution of all craving and desire ratings was inspected to visualize differences across tasks (alcohol vs. neutral image viewing) and states (typical vs. abstained conditions). To evaluate differences in ratings across tasks and states, a hierarchical mixed-effects regression model was used to examine potential interactions. Ratings, imagery task, and drinking state were included in the model as follows: Rating: 1=Desire, 2=Crave; Task: 1=Neutral, 2=Alcohol; State 1=Typical, 2=Abstinence. Wilks’ Lambda statistics were examined.

## RESULTS

Demographic information can be found in Table 1. Thirty-nine participants (22 female) completed the full study protocol. Participants averaged 41 years-of-age and were regular alcohol consumers for an average of nearly 25 years. Overall, participants consumed an average of 17 (±5) drinks per week or 2.9 (±0.9) drinks per day, with males consuming [23 (±6) per week or 3.7 (±1.2) per day] significantly more than females [13 (±5) per week or 2.4 (±0.7) per day; t = 5.22, p < 0.001]. Race, age, and total years drinking were not significantly different between males and females. Average scores from alcohol related surveys administered during the screening or baseline visits are included in Table 2 (Alcohol Craving Experience (ACE) Questionnaire (Statham et al., 2011) and the Alcohol Use Disorder Identification Test (AUDIT) (WHO, 2001). Table 2 also includes correlation values between total average scores and ratings of desire and craving. Overall, ACE scores were moderately correlated with ratings of desire and craving, while total number of drinks consumed in the past 90 days was not significantly correlated with desire or craving scores, overall. Interestingly, AUDIT scores were only significantly correlated with ratings of craving following neutral imagery viewing, following typical drinking and abstinence.

**Table 1.** Participant demographics. Displayed as N (%) or Mean (±SD). Average consumption was measured as number of standard drinks (containing approximately 14g of alcohol, e.g. 12oz of beer, 5oz of wine, or 1.5oz of distilled spirits). * significantly different at *p* < 0.001, tested via t-test.

| Demographic | Overall | Male | Female |
| --- | --- | --- | --- |
| N | 39 | 17 (44%) | 22 (56%) |
| White | 35 (90%) | 15 (88%) | 20 (91%) |
| African American or Black | 4 (10%) | 2 (12%) | 2 (9%) |
| Age | 41.2 (±11.7) | 38.8 (±9.8) | 43 (±13) |
| Total Years Drinking | 24.7 (±11.8) | 21.9 (±9.2) | 26.9 (±13.2) |
| Average consumption per week | 17 (±5) | 23 (±6)* | 13 (±5)* |
| Average consumption per day | 2.9 (±0.9) | 3.7 (±1.2)* | 2.4 (±0.7)* |

**Table 2.** Alcohol related survey responses and correlation with ratings, collected in the MRI scanner across drinking states (typical drinking and abstinence), imagery task (neutral or alcohol), and rating type (desire or crave). Total scores on the ACE were out of 110 and on the AUDIT out of 50. Significant relationships are bolded.

|  | Alcohol Craving Experience (ACE) Questionnaire | Alcohol Use Disorder Identification Test (AUDIT) | Total Drinks Consumed in the Previous 90 Days |
| --- | --- | --- | --- |
| Overall Average Totals | 13.54 (±10.00) | 18.41 (±2.92) | 224.10 (±91.25) |
| Typical Neutral Desire | **$r = 0.422, p = 0.020$** | $r = 0.357, p = 0.053$ | $r = 0.007, p = 0.969$ |
| Typical Neutral Crave | **$r = 0.713, p < 0.001$** | **$r = 0.479, p = 0.007$** | $r = 0.021, p = 0.911$ |
| Typical Alcohol Desire | **$r = 0.454, p = 0.012$** | $r = 0.349, p = 0.058$ | $r = 0.100, p = 0.599$ |
| Typical Alcohol Crave | **$r = 0.601, p < 0.001$** | $r = 0.354, p = 0.055$ | $r = 0.023, p = 0.906$ |
| Abstain Neutral Desire | **$r = 0.362, p = 0.049$** | $r = 0.357, p = 0.061$ | $r = 0.313, p = 0.092$ |
| Abstain Neutral Crave | **$r = 0.512, p = 0.004$** | **$r = 0.385, p = 0.036$** | $r = 0.264, p = 0.159$ |
| Abstain Alcohol Desire | $r = 0.198, p = 0.294$ | $r = 0.294, p = 0.115$ | $r = 0.172, p = 0.364$ |
| Abstain Alcohol Crave | $r = 0.351, p = 0.057$ | $r = 0.281, p = 0.133$ | $r = 0.117, p = 0.538$ |

Descriptive statistics of EMA responses are included in Table 3 and Figure 2. These data depict desire and craving scores reported across each day of the two experimental sessions, categorized as occurring pre-drinking, post-drinking, or during abstinence. Desire scores were consistently different from and greater than craving scores (*t* = 21.783, *p* < 0.001). Desire scores appear to increase from pre-drinking to post-drinking, with desire ratings collected during abstinence exceeding ratings collected pre-drinking (although lower than post-drinking). Although both variables showed similar patterns of response, the average values of desire ratings were consistently higher than average craving ratings (*t* = 7.275, *p* < 0.001). Predicted craving and desire ratings across the day, generated from the EMA hierarchical modeling, are included in Figures 3 and 4. This modeling showed that craving ratings increased throughout the day (γ = 9.335, SE = 0.883, *p* < 0.001) with a quadratic decrease in the late afternoon and evening (γ = −1.329, SE = 0.152, *p* < 0.001). Although pre-drinking and abstinence ratings of craving were similar, craving scores increased significantly following drinking (γ = 30.475, SE = 11.342, *p* = 0.007). Desire ratings also increased across the day (γ = 15.531, SE = 1.162, *p* < 0.001) with a late day decrease (γ = −2.592, SE = 0.120, *p* < 0.001). Compared to days of abstinence, desire ratings were lower prior to drinking (γ = −30.421, SE = 12.775, *p* = 0.017) and higher following drinking (γ = 61.685, SE = 14.925, *p* < 0.001). Full values from the statistical model are included in Tables 4 and 5. The ICC for craving ratings was 0.2633 and for desire ratings was 0.1564, indicating approximately 26% of the variance in craving scores was between subjects whereas about 74% was within subject, and approximately 16% of the variance in desire ratings was between subjects leaving 84% within subject, respectively. Despite these differences in within-person variability, the variables were strongly correlated (*r* = 0.76, *p* < 0.001), as were the craving and desire standard deviations (*r* = 0.54, *p* < 0.001).

**Figure 2.**
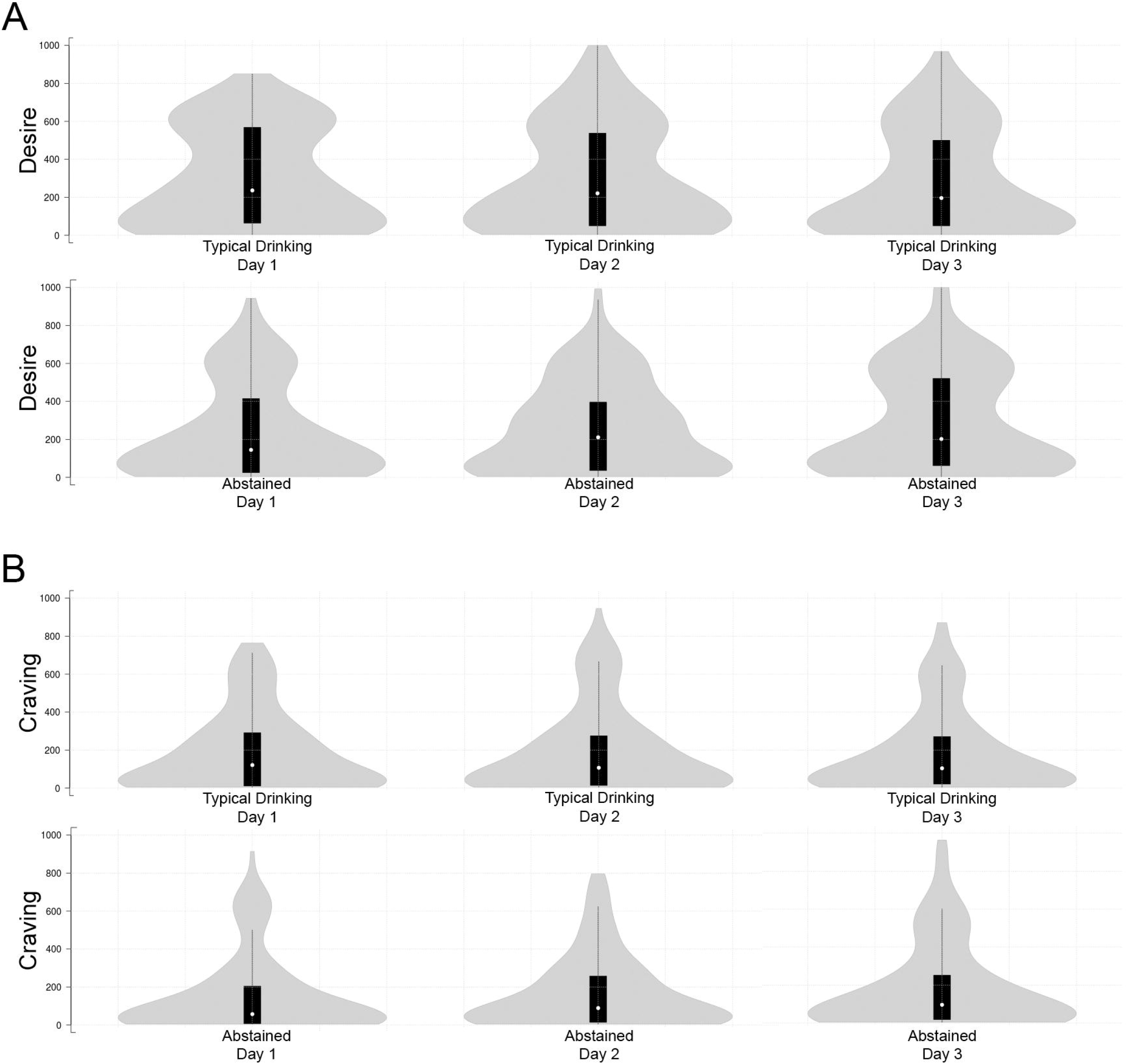
Desire and craving ratings collected during the EMA protocol. In the plots, white circles show the medians; black box limits indicate the 25^th^ and 75^th^ percentiles; whiskers extend 1.5 times the interquartile range; and grey polygons represent density estimates of data and extend to extreme values. See Table 2 for descriptive statistics.

**Figure 3.**
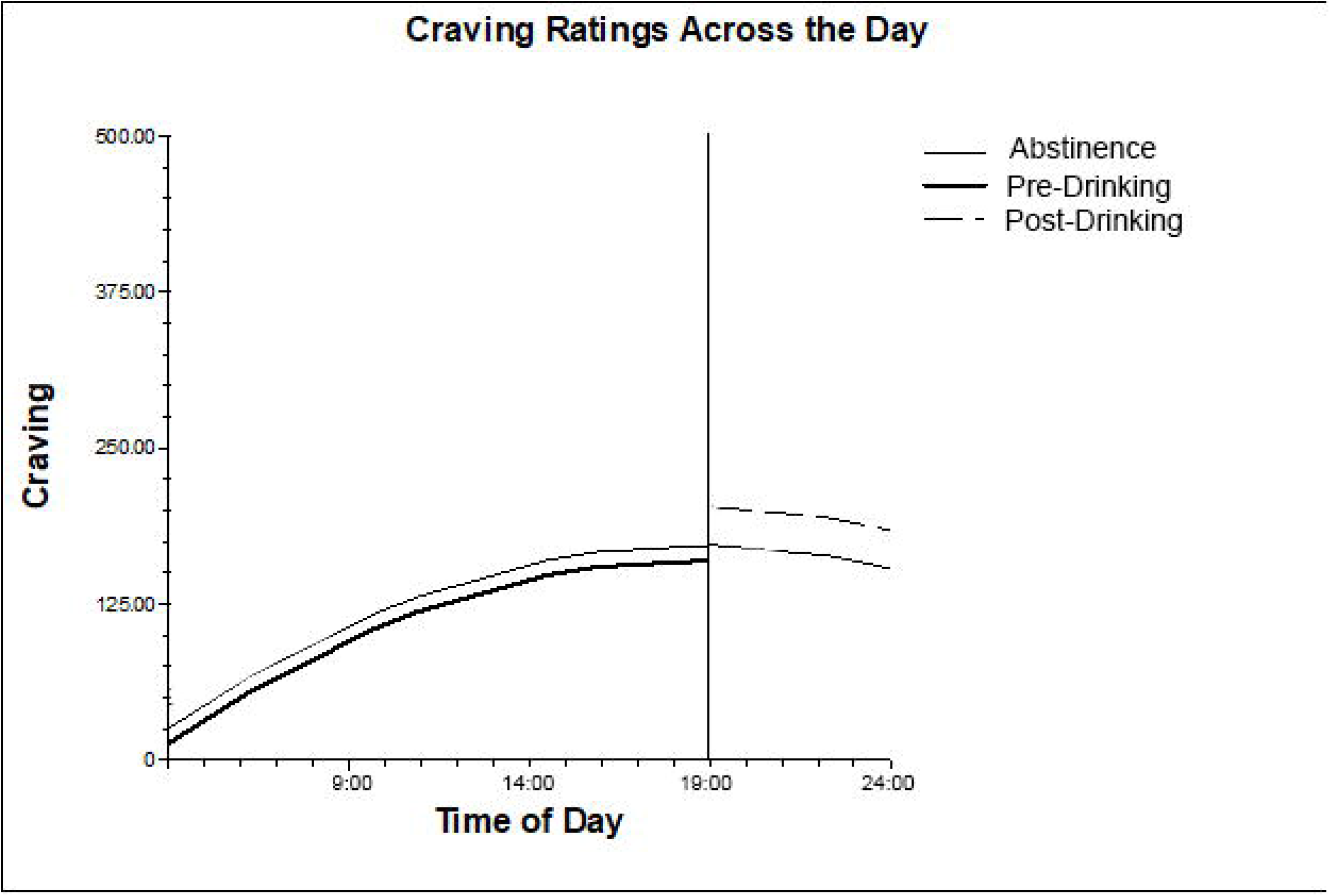
Predictive modeling of craving ratings from EMA responses, depicted across the day, split as pre-drinking, post-drinking (from the participant’s typical drinking days), and abstained. The x-axis depicts time of day and the y-axis shows craving scores. The vertical line represents time of consumption on typical drinking days. See Table 2 for descriptive statistics of EMA responses.

**Figure 4.**
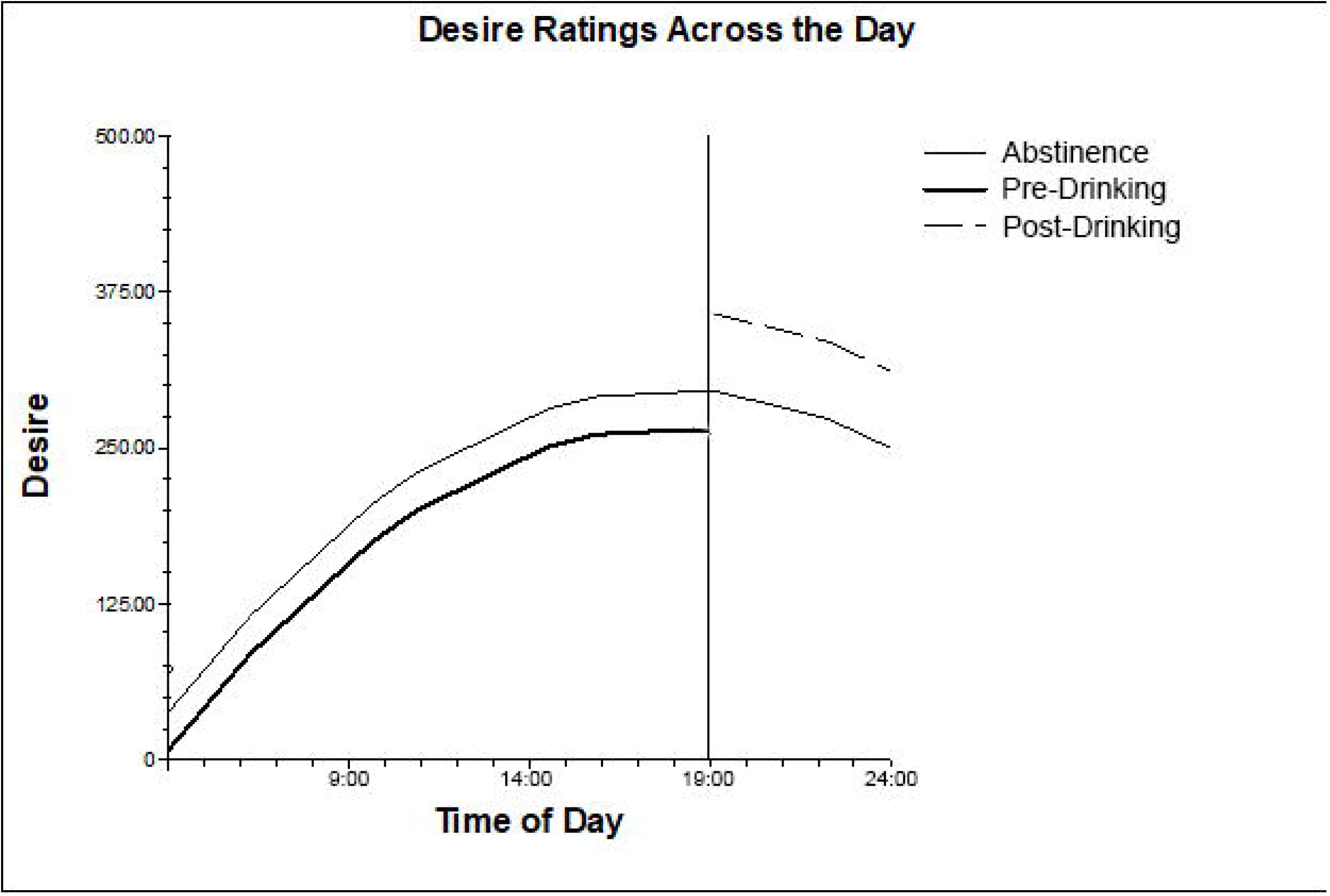
Predictive modeling of desire ratings from EMA responses, depicted across the day, split as pre-drinking, post-drinking (from typical drinking days), and abstained. The x-axis depicts time of day, and the y-axis shows desire scores. The vertical line represents time of consumption on typical drinking days. See Table 2 for descriptive statistics of EMA responses.

**Table 3.** EMA desire and craving ratings collected during the 3-day experimental periods. Scores are displayed as Mean (±SD) and (Range). Potential ratings ranged 0 – 1000. Responses are categorized as pre-drinking or post-drinking during the 3 days of typical drinking or during the period of abstinence.

| Rating Scale/Experimental Period | Day 1 | Day 2 | Day 3 | 3-Day Average |
| --- | --- | --- | --- | --- |
| **Desire** |  |  |  |  |
| Pre-Drinking | 128.97 (±115.31)<br>(0 – 396) | 134.82 (±129.76)<br>(0 – 500) | 162.78 (±134.93)<br>(0 – 478.50) | 142.17 (126.67)<br>(0 – 500) |
| Post-Drinking | 312.87 (±160.28)<br>(69.75 – 804) | 305.02 (±176.56)<br>(58.33 – 671.33) | 327.53 (±146.04)<br>(144 – 751.83) | 315.14 (160.96)<br>(58.33 – 804) |
| Abstinence | 162.09 ($\pm 117.11$)<br>(0 – 439.14) | 187.79 ($\pm 143.21$)<br>(0.67 – 567.67) | 219.89 ($\pm 145.13$)<br>(21.50 – 669.50) | *189.92 (135.15)*<br>(0 – 669.50) |
| **Craving** |  |  |  |  |
| Pre-Drinking | 72.89 ($\pm 101.34$)<br>(0 – 414) | 77.20 ($\pm 102.19$)<br>(0 – 500) | 106.09 ($\pm 124.63$)<br>(0 – 464) | *85.39 (109.39)*<br>(0 – 500) |
| Post-Drinking | 184.45 ($\pm 184.94$)<br>(0 – 743) | 174.13 ($\pm 180.94$)<br>(0 – 646.50) | 167.21 ($\pm 140.35$)<br>(0 – 490.33) | *175.26 (168.74)*<br>(0 – 743) |
| Abstinence | 83.13 ($\pm 87.34$)<br>(0 – 337.50) | 115.01 ($\pm 130.27$)<br>(0 – 443) | 124.69 ($\pm 122.48$)<br>(0 – 454.33) | *107.61 (113.36)*<br>(0 – 454.33) |

**Table 4.** Results from hierarchical modeling of craving EMA ratings. Significant effects are bolded.

| Effect | Gamma | Standard Error | p-value |
| --- | --- | --- | --- |
| Intercept | **155.76** | **18.55** | **< 0.001** |
| Time of Day – Linear Slope | **9.34** | **0.88** | **< 0.001** |
| Time of Day – Curvilinear Slope | **-1.33** | **0.15** | **< 0.001** |
| Pre-Drinking Slope | -12.02 | 9.70 | 0.216 |
| Post-Drinking Slope | **30.48** | **11.34** | **0.007** |

**Table 5.** Results from the hierarchical modeling of desire EMA ratings. Significant effects are bolded.

| Effect | Gamma | Standard Error | p-value |
| --- | --- | --- | --- |
| Intercept | **274.10** | **19.13** | **< 0.001** |
| Time of Day – Linear Slope | **15.53** | **1.16** | **< 0.001** |
| Time of Day – Curvilinear Slope | **-2.59** | **0.20** | **< 0.001** |
| Pre-Drinking Slope | **-30.42** | **12.78** | **0.017** |
| Post-Drinking Slope | **61.69** | **14.93** | **< 0.001** |

In an effort to examine potential patterns obscured by the within subject variability, predictive modeling was used to determine whether craving ratings were predicted by desire ratings, while controlling for linear and quadratic times of day, as well as pre- and post-drinking differentiation. Time of day and pre- versus post-drinking did not significantly predict craving ratings, while desire ratings show a significant relationship with craving scores (γ = 0.508, SE = 0.013, *p* < 0.001). To visualize overall differences between desire and craving EMA ratings, raw craving scores were subtracted from desire scores across all EMA responses from all participants (Figure 5). The resulting differences scores depicted overall differences in ratings: positive differences indicate greater desire ratings, and negative difference scores indicates greater craving ratings. If a systematic difference between desire and craving ratings is present, a fairly flat distribution of difference scores would be observed. However, the waterfall plot shows the wide variability in the difference between ratings between subjects, but does not show a consistent trend in difference scores, indicating no global pattern of difference. However, following abstinence, a large proportion of participants appear to rate their desire much higher than their craving, and during typical consumption, while many participant maintain this “desire > craving” pattern of response, a subset of participants appear to be shifting toward a smaller or zero difference between ratings.

**Figure 5.**
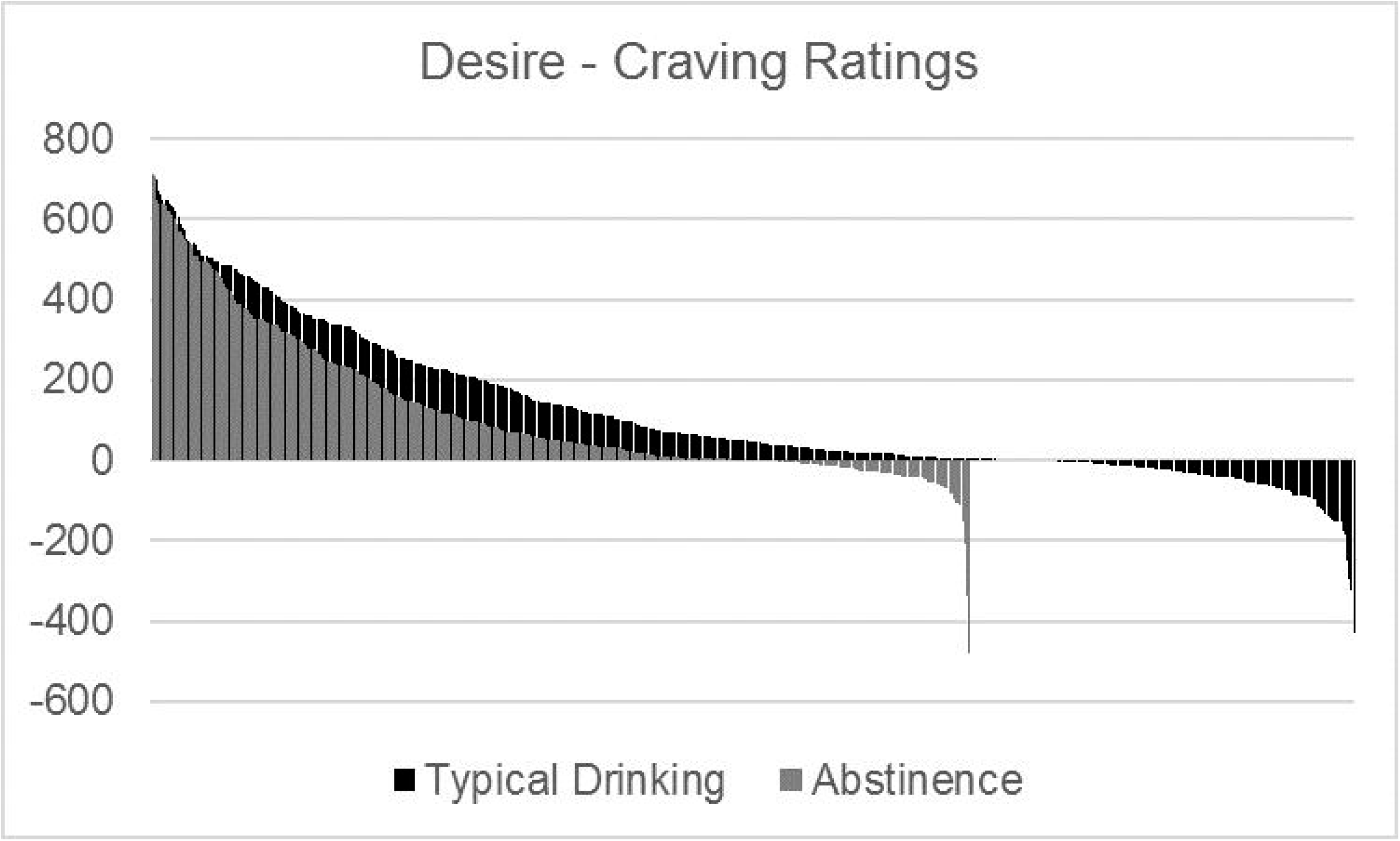
EMA responses, desire ratings minus craving ratings, separated by drinking state. Positive difference scores indicate a greater desire rating, while a negative difference score indicates a greater craving rating. Difference scores across all participants and surveys sorted by magnitude.

Average ratings collected following the imagery tasks across both experimental drinking conditions are included in Table 6 and Figure 6. These values show higher desire ratings than craving ratings across both tasks and conditions, increases to both desire and craving when viewing the alcohol related images compared to the neutral images, and (visually present but not statistically significant) increases following abstinence compared to typical drinking. None of the in-scanner ratings (across tasks, drinking states, and rating type) were significantly associated with levels of alcohol consumption, assessed via the TLFB. Full results from this statistical model are included in Table 7. The mixed-model regression used to evaluate these differences showed a significant effect of rating type (indicating a significant difference overall between desire and craving ratings; *F* = 44.20, *p* < 0.0001) as well as a significant interaction between rating type and task (*F* = 11.52, *p* = 0.0003), indicating the difference between rating types varies between the two imagery tasks. However, no significant interactions including drinking state were found, indicating no significant difference in ratings collected following the MRI imagery scans between typical drinking and abstinence. Of note, an additional significant interaction was found between ratings and participants’ age (*F* = 4.62, *p* = 0.0197) such that ratings increased as age increased.

**Figure 6.**
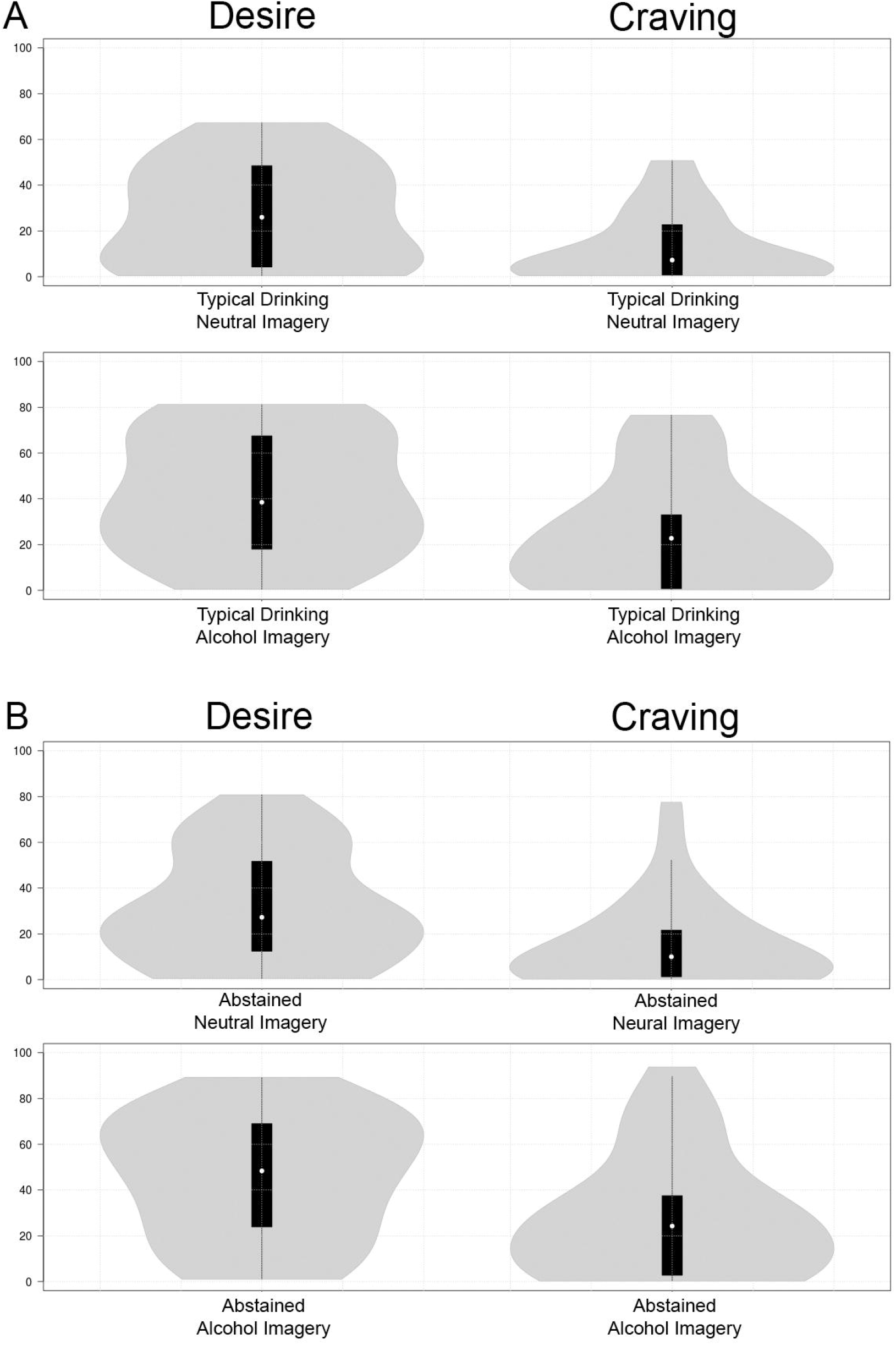
Desire and craving ratings collected following the imagery MRI scans. In the plots, white circles show the medians; black box limits indicate the 25^th^ and 75^th^ percentiles; whiskers extend 1.5 times the interquartile range; and grey polygons represent density estimates of data and extend to extreme values. Ratings of desire were significantly greater than ratings of craving, and ratings increased significantly following the alcohol imagery compared to the neutral imagery. Although there appears to be an increase in ratings across state, this effect was not significant.

**Table 6.** Average desire and craving ratings collected following the functional MRI scans (neutral imagery, alcohol imagery) during the typical drinking scan and the abstinence scan. Scores are displayed as Mean (±SD) and (Range).

| State/Task | Desire | Crave |
| --- | --- | --- |
| **Typical Drinking** |  |  |
| Neutral Imagery | 27.29 ($\pm 22.32$)<br>(0.50 - 67.25) | 13.39 ($\pm 15.27$)<br>(0.50 - 50.75) |
| Alcohol Imagery | 42.55 ($\pm 25.92$)<br>(0.50 - 81.25) | 24.82 ($\pm 24.15$)<br>(0.25 - 76.50) |
| **Abstained** |  |  |
| Neutral Imagery | 31.10 ($\pm 24.21$)<br>(0.50 - 80.75) | 15.59 ($\pm 19.33$)<br>(0.25 - 77.50) |
| Alcohol Imagery | 46.16 ($\pm 26.83$)<br>(1.00 - 89.25) | 27.66 ($\pm 26.04$)<br>(0.25 - 93.75) |

**Table 7.** Results from hierarchical modeling of ratings collected in the scanner. Wilks’ Lambda statistics are shown. Significant and trending effects and interactions are bolded. (Rating: 1=Desire, 2=Crave; Task: 1=Neutral, 2=Alcohol; State: 1=Typical, 2=Abstinence)

| Effect | Beta | F Value | p-value |
| --- | --- | --- | --- |
| **Rating** | **0.2205** | **44.2** | **< 0.0001** |
| **Rating\*Task** | **0.5205** | **11.52** | **0.0003** |
| Rating*State | 0.9945 | 0.07 | 0.9338 |
| Rating*Task*State | 0.9957 | 0.05 | 0.9471 |
| **Rating\*Sex** | **0.8169** | **2.80** | **0.0799** |
| **Rating\*Age** | **0.7304** | **4.62** | **0.0197** |
| Rating*Total Drink Count | 0.8909 | 1.53 | 0.2360 |
| Rating*Task*Sex | 0.0727 | 0.98 | 0.3894 |
| Rating*Task*Age | 0.9273 | 0.98 | 0.3894 |
| Rating*Task*Total Drink Count | 0.9861 | 0.18 | 0.8393 |
| Rating*State*Sex | 0.9378 | 0.83 | 0.4484 |
| Rating*State*Age | 0.9819 | 0.23 | 0.7959 |
| Rating*State*Total Drink Count | 0.9725 | 0.35 | 0.7058 |
| Rating*Task*State*Sex | 0.9711 | 0.37 | 0.6931 |
| Rating*Task*State*Age | 0.9355 | 0.86 | 0.4344 |
| Rating*Task*State*Total Drink Count | 0.9895 | 0.13 | 0.8769 |

## DISCUSSION

Among non-binging regular drinkers, ratings of desire and craving for alcohol are consistently different while drinking according to a person’s typical routine or abstaining, throughout the day, and when viewing alcohol cue imagery. Although these results cannot definitively indicate that participants do not conceptualize desire and craving as synonymous, this study’s key finding is that the ecologically valid ratings collected across the day and ratings collected following the alcohol imagery MRI scan do clearly demonstrate a distinction between self-reported experiences of craving and desire for alcohol. Although the intraclass correlations are merely a descriptive finding, they suggest desire ratings may be more labile in this sample, compared to ratings of craving. Contrary to expectations, ratings within each category (i.e. desire or craving) did not significantly differ across drinking states (typical drinking and abstinence), although the observed differences between craving and desire ratings were maintained across both pre- or post-drinking states. Additionally, both craving and desire ratings increased following alcohol cue exposure. Although many researchers have found that drinkers diagnosed with AUD do not synonymously report craving and desire experiences (W.H.O., 1955, Kozlowski et al., 1989, Breese et al., 2006, Gordon et al., 2006, Miller and Gold, 2011, Tuithof et al., 2014), to our knowledge, this is the first study to demonstrate that participants who do not experience self-reported behavioral issues associated with their consumption also significantly differ in their ratings of their craving and desire to drink. The drinking population sampled for this study does represent a large portion of the riskier drinking population in the US (NIAAA, 2011, NSDUH, 2019). Further studies are necessary to determine how these results extrapolate to other populations, but the differences found between ratings in this study present a reasonable foundation for the continuation of this line of research.

Across all measures, desire ratings, on average, were greater than craving ratings. However, this difference was not systematically distributed (e.g. if desire had been consistently rated as twice the value craving), suggesting a nuanced semantic difference between the psychological and physiological states drinkers classify as “craving” and “desire”. This is clearly demonstrated in Figure 5, where we see some participants maintain their much greater desire rating across drinking states, while other appear to rate the scales more similarly (difference closer to 0) during normal drinking. Because of these findings, we do not feel a simple explanation such as “craving” is typically interpreted as a stronger version of “desire” is likely to capture the full range of potentially subtle variations in ratings. Even when controlling for age, sex, and total number of drinks consumed in the past 90-days (Table 6), no overall pattern of difference in ratings was observed, with significant (and trending) differences by age and sex found only when examining the main effect of ratings.

As alcohol research continues to advance toward preventive therapies and more effective treatments for AUD, a more thorough understanding of the craving experience and well defined operational definitions when asking participants to report a specific psychological or physiological experience could fill current research gaps or expand understanding of existing results. The long-term hope is to accurately capture “craving” (as differentiated from “desire”) as a risk factor for developing AUD, an important symptom when assessing and diagnosing AUDs, as an indicator of relapse risk, and as a motivator for the drinker seeking clinical assessment. Because the concept of craving plays such a dominant role in alcohol-related medicine and research (Sullivan et al., 1989b, Miller and Gold, 1994, Gordon et al., 2006, APA, 2013), alcohol researchers must know they are creating and executing experiments which probe and effect of the same physiological and psychological experience. Perhaps most importantly, equipped with the knowledge that “craving” and “desire” may not have widely agreed upon meanings to a lay audience, addiction specialists, and researchers should strongly consider clear and definitive operational definitions for the states they plan to examine to be sure all participants or patients are reporting based on the same experience.

This investigation was not without limitations. Primarily, this study has a limited sample size which needs to be expanded into larger samples. Additionally, this small sample only included non-binging regular alcohol consumers, and as such these results cannot be immediately extrapolated into different drinking populations. This study should be replicated in a broader collection of alcohol consumers, ranging from social drinkers to treatment seeking patients diagnosed with an AUD. Finally, all questions probing craving and desire were single items, as opposed to the more complex instruments traditionally used to probe craving. A more complete battery assessing urge to drink and alcohol seeking could clarify the findings of this study. However, even considering these limitations, we would like to emphasize that this approach has several strengths, including ecological validity and reduced recall bias, as well as the inclusion of EMA data and rating data collected after viewing neutral and alcohol images.

In summary, the concepts of craving and desire are not well defined in alcohol or other substance use research, and researchers and participants may conceptualize and rate these terms differently. In the healthy, non-binging regular alcohol consumers studied in this protocol, self-reported ratings of desire and craving for alcohol appeared to capture disparate experiences. However, future studies are needed to explore the differences between these biological and conceptual differences, throughout the continuum of drinking populations.

## Funding

NIAAA P50 AA026117

NIAAA P01 AA02099

NIAAA T32 AA007565

All Authors confirm they have no conflicts of interest to declare.

